# AlphaFold 3 accurately models natural variants of Helicobacter pylori catalase KatA

**DOI:** 10.1101/2025.06.02.657526

**Authors:** Arden Baylink

## Abstract

Subtle changes in protein sequence can equate to large changes in function, such as enabling pathogens to evade the immune system, hindering antibody recognition of antigens, or conferring antibiotic resistance. Even single amino acid substitutions may alter ligand binding affinity, enzymatic activity, and protein stability. Yet, due to limitations in time and resources, proteins closely related in sequence to those already characterized often remain unexamined. AlphaFold has emerged as a promising tool for protein structure prediction, though its utility in modeling single amino acid substitutions remains uncertain. In this study, we assessed AlphaFold 3’s accuracy in modeling natural variants of the *Helicobacter pylori* catalase KatA by comparing its predictions to a novel high-resolution crystal structure of KatA from strain SS1. This variant contains key substitutions at residues 234, 237, 255, and 421 relative to the well-characterized strain 26695. AlphaFold 3 models accurately reproduced the global structure and local conformations of most variant residues, with high fidelity in conservative substitutions but variable accuracy in more flexible or interface-exposed sites. We further explored how user inputs, such as incorrect oligomeric states or sequence modifications, influence prediction quality. While AlphaFold 3 consistently produced high-quality models, deviations at variant sites occurred when incorrect oligomeric states were specified. Our findings highlight both the strengths and limitations of AlphaFold 3 in modeling natural protein variants and underscore the importance of accurate user input for reliable structural predictions.

**Importance:** Experimental structure determination is rarely performed for natural protein variants possessing only minor amino acid differences from published structures, even though small substitutions can significantly impact structure and function. Here, we present a case study showing that AlphaFold 3 can accurately model the structures of natural protein variants. However, providing an incorrect oligomeric state can reduce model accuracy—an error that non-expert users may easily make.

## Introduction

Proteins in nature exist as a spectrum of amino acid variants ^1–4^. These sequence differences may result from genetic drift, which over evolutionary timescales can lead to gain- or loss-of-function events ^5^. In some cases, however, single amino acid changes can immediately influence protein function by altering ligand-binding specificity, enzymatic efficiency, conformational dynamics, thermostability, immunogenicity, or other properties ^6–9^. Amino acid variation underlies positive selection in host-pathogen evolutionary relationships and antimicrobial resistance ^10–12^. The structural and functional consequences of such amino acid changes can be studied and understood using experimental structure-determination methods like protein crystallography, which can reveal unexpected alterations not readily predicted by sequence analysis alone ^13,14^. However, structural studies of proteins similar in sequence, such as natural protein variants, are not commonly undertaken due to constraints in time and resources, limiting our understanding of how natural variances affect structure and function.

AlphaFold has emerged as a promising tool to address some of these challenges ^4,15,16^. Widely recognized as a significant advancement in protein structure prediction, AlphaFold has been used successfully to model new protein structures ahead of experimental validation, and is continually being updated and developed with improved features ^15–17^. There is disagreement in the literature about the application of AlphaFold modeling of single amino acid substitutions. Studies suggest that AlphaFold’s predictive accuracy is system-dependent and heavily influenced by the quality of experimental data used during its training ^18,19^. The developers explicitly caution against using AlphaFold to predict the structural impact of single amino acid substitutions, such as destabilizing mutations (https://alphafold.ebi.ac.uk/faq) ^20^. A recent study showed that AlphaFold’s predictions for the energetic consequences of single mutations poorly correlate with experimentally determined protein stabilities ^9,20^. However, new ways to circumvent some of these limitations have been reported ^4^. As a result, the utility of AlphaFold in accurately modeling local structural variations remains uncertain.

Despite these limitations, AlphaFold has significantly broadened access to protein structure prediction, enabling researchers without structural biology expertise to model proteins and explore mechanistic hypotheses ^21^. The online AlphaFold 3 server, for example, only requires users to input a primary amino acid sequence and oligomerization state and returns a model within minutes ^22^. However, this ease of access also introduces risks. Users unfamiliar with structural biology may provide suboptimal input—such as incorrect oligomeric states—or misinterpret the resulting models, potentially compromising prediction accuracy. Thus, user input represents another variable affecting AlphaFold’s performance.

In this study, we evaluated the ability of AlphaFold 3 to model previously uncharacterized natural variants of the catalase KatA from the gastric pathogen *Helicobacter pylori*. This system is well-suited as a case study because *H. pylori* has a notoriously high mutation rate, leading to numerous naturally occurring protein variants that may contribute to its pathogenicity and resistance to antibacterial drugs ^23,24^. High-resolution crystal structures exist for KatA, though they are limited to a single strain (26695), providing AlphaFold with accurate experimental data as a basis for modeling ^25,26^. We performed a straightforward test by first solving a novel KatA crystal structure from strain SS1, which contains several sequence variations from the published crystal structures, and then assessed the accuracy of predictions made by the Alphafold 3 server ^22^. We used the AlphaFold 3 server under conditions that mimic how a non-structural biologist might approach modeling, and tested how slightly incorrect inputs impact the resulting models. This study serves as a test case for AlphaFold’s capability to predict the structures of naturally occurring variants and offers insights into how user-provided input may influence modeling accuracy.

## Results & Discussion

### Characterization of natural KatA variants

KatA is the only catalase produced by *H. pylori*, is monofunctional, and, like other catalases, converts hydrogen peroxide (H_2_O_2_) to water and oxygen through a heme b prosthetic group, thereby protecting the bacterium against oxidative stress ^25–27^. The enzyme is 505 amino acids in length and forms a tetramer, with one active site per monomer and solvent channels that enable the buried heme groups to access and dismutate H_2_O_2_ substrate (Fig. 1A) ^25^. To our knowledge, no prior work has evaluated the natural variants of *H. pylori* KatA, and so we conducted searches of the published *H. pylori* genomes for KatA orthologues, retrieving 1,931 sequences, with 1,922 being unique (Data S1).

**Fig. 1.**
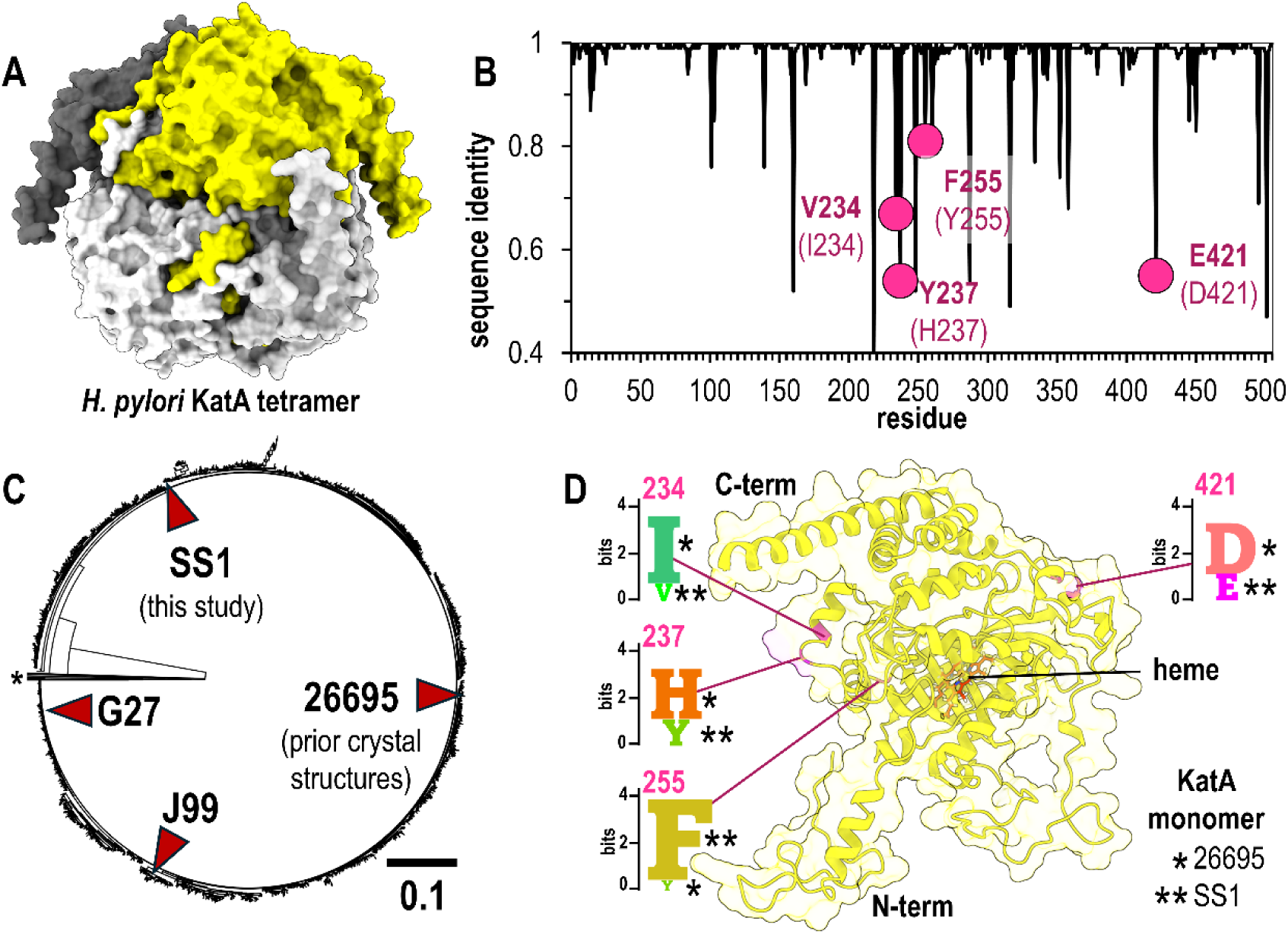
Natural variants of *H. pylori* KatA. A. The biologically-relevant KatA tetramer, colored by chain (white, light gray, dark gray, yellow). B. Sequence identity at each amino acid position over 1,931 *H. pylori* KatA sequences. Natural variants of interest in this study highlighted in pink, with the residue for SS1 strain in bold and corresponding residue for strain 26695 in parentheses. C. Relatedness tree for *H. pylori* KatA sequences with commonly-used model strains noted. A small number of divergent sequences, which could potentially be misannotations from other species, are not shown (*). D. The position within the *H. pylori* KatA monomer for the variants of this study are indicated in pink, along with SeqLogo plots showing conservation patterns at each site across all identified sequences. Single Asterix (*) indicates the residue present in strain 26695 and double Asterix (**) indicates the residue present in strain SS1.

To compare variation at each amino acid site, we performed multi-sequence alignment and assessed sequence conservation at each position. While KatA is overall highly conserved, approximately 20 sites show less than 90% sequence identity (Fig. 1B). Among these, we discovered that residues 234, 237, 255, and 421 are common sites of variation, which for the previously-characterized KatA from strain 26695 are Ile, His, Tyr, and Asp, respectively (Fig. 1B) ^25^. Position 234 and 237 constitute part of the solvent channel opening, 255 is solvent exposed and makes no other notable interactions, and 421 is involved in the stabilizing tetrameric interactions with the N-terminus of another partner chain (PDB: 1qwl, 1qwm,2a9e, 2iqf) ^25,26^.

Strain 26695 is a clinical isolate used as a research model, as are the strains G27, J99, and SS1, the latter being a mouse-adapted strain commonly employed in animal infection studies ^28–33^. To visualize sequence variation of KatA across *H. pylori* strains, we generated a relatedness tree for KatA variants, and noted that these aforementioned strains apparently represent different clades (Fig. 1C). Examining the variation across all *H. pylori* strains at the four sites mentioned above, shows each to be dominated by two different amino acids; 234 (Ile/Val), 237 (His/Tyr), 255 (Phe/Tyr), and 421 (Asp/Glu) (Fig. 1D). Interestingly, we noted that while the earlier KatA 26695 crystal structures represent the subpopulation of Ile, His, Tyr, and Asp at these sites, the SS1 strain possesses the complementary Val, Tyr, Phe, and Glu (Fig. 1D). Hence, we decided to pursue structural studies of KatA SS1 to obtain the first experimental characterization of the structure of these variants; we reasoned this also would provide us with a small but straightforward set of standards for Alphafold 3 modeling spanning a few different residue types and locations across the protein (Fig. 1D).

### Structure of H. pylori strain SS1 KatA positions of variant residues

The crystal structure of KatA from strain SS1, which we henceforth refer to as KatA_SS1_, was solved at 1.87 Å resolution in space group P2_1_ 2 2_1_ with two chains in the asymmetric unit, i.e. one half of the biologically-relevant tetramer. This was serendipitous since this novel structure represents a new crystal form with different crystal packing interactions than previously- published structures ^25,26^, and the two non-crystallographic symmetry-related chains offer two separate views for the amino acids of interest. The electron density was clear and easy to interpret for the majority of the structure, and only portions of the N- and C-terminus were not modeled (see Methods). An overlay of the two chains of KatA_SS1_ with the four structures from strain 26695 (KatA_26695_) showed high similarity, as expected, with no major differences in global architecture (Fig. 2A).

**Fig. 2.**
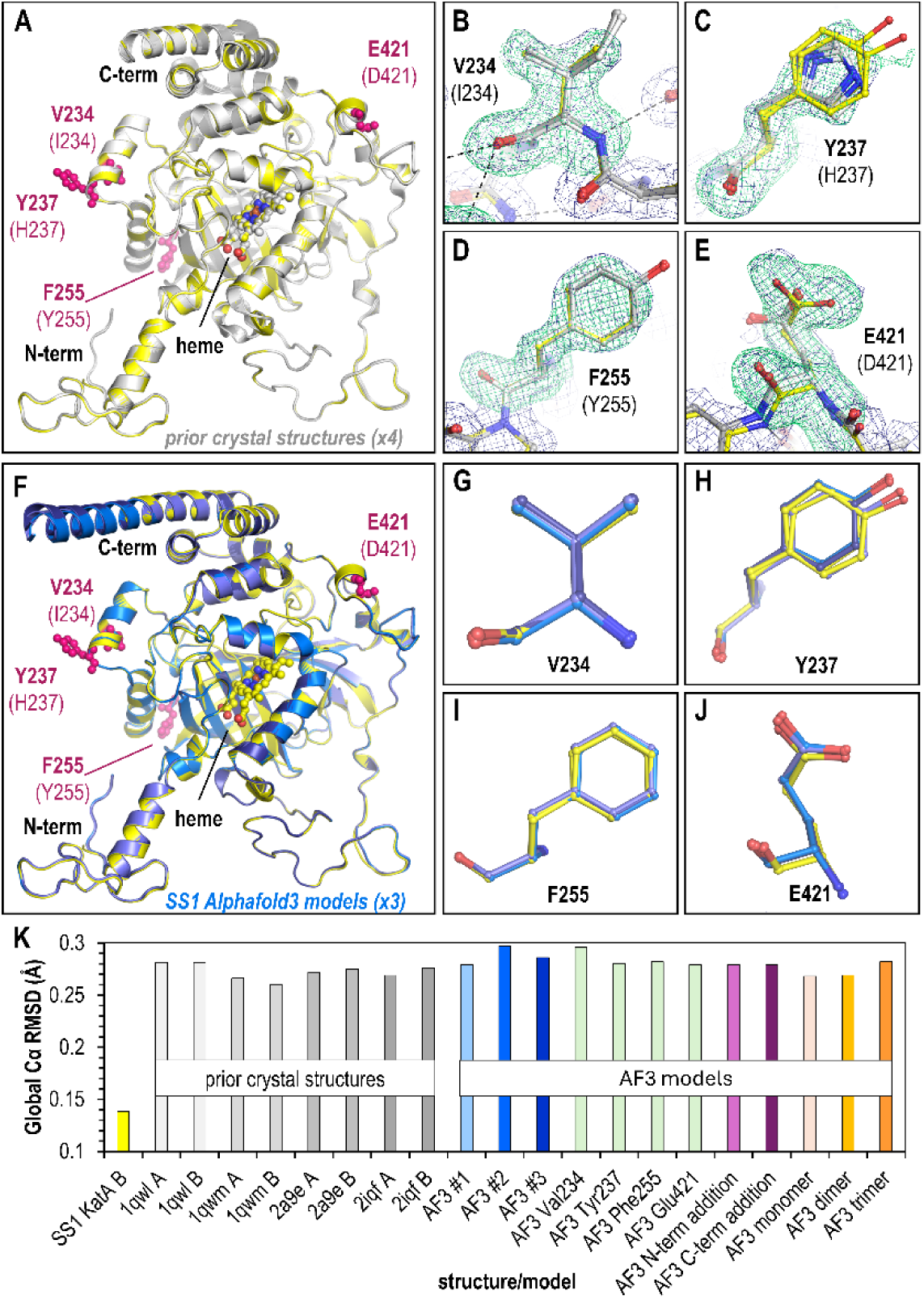
Comparisons of crystal structure and Alphafold 3 models of KatA_SS1_. A. Overlay of crystal structure KatA_SS1_ (2 chains, yellow), and crystal structures of KatA_26659_ (light gray, 4 chains). Variant sites are noted in pink. B-E. Poses of variant residues within KatA_SS1_, as indicated. KatA_SS1_ crystal structure chains are shown in yellow and KatA_26995_ crystal structure chains in light gray. Green mesh is Fo-Fc omit map electron density at 3.5 σ averaged over the two non-crystallographic symmetry (NCS) chains, and dark blue mesh is 2Fo-Fc NCS-averaged electron density at 1.0 σ of the final model. F. Overlay of the KatA_SS1_ crystal structure (yellow) with three Alphafold 3 models (light blue, blue, dark blue) using native sequence and tetramer oligomerization as input parameters. G-J. Overlay of each variant residue comparing crystal structure (yellow) positions to Alphafold 3 predictions (blue). K. Global root-mean-square-deviation (RMSD) values for each structure and model in relation to crystal structure KatA_SS1_ chain A.

To assess the positions of the variant sites, we deleted positions 234, 237, 255, and 421 to generate an omit map, which reduces modeling bias in the electron density, and also calculated non-crystallographic symmetry maps averaged across the two chains in the asymmetric unit (Fig. 2B-E). For residue 234 (Ile234 in KatA_26695_), the density can be clearly interpreted as a Val, positioned similar to Ile234 without the C_δ_ (Fig. 2B). For Tyr237, the density is much weaker, and small difference density peaks near C_β_ suggest it may adopt multiple conformations. The dominant residue position is well-defined through C_β_, and the general plane of the phenol ring is apparent, but the side chain O is not resolved (Fig. 2C). The Tyr237 residues in the two KatA_SS1_ chains also adopt slightly different positions (Fig. 2C). The electron density is unambiguous for both residue 255, which can be readily modeled as a Phe, and for residue 421, a Glu, with all atoms of these residues clearly visible in the electron density (Fig. 2D, Fig. 2E). Overall, the positions of these variants in the KatA_SS1_ structure are well-supported by the electron density and adopt conformations that could be reasonably predicted by an experienced structural biologist.

### Alphafold 3 accurately predicts KatA_SS1_ variant poses

Having experimentally-determined the positions of the four KatA_SS1_ variant sites, for which no previous structure has been reported, and therefore are not part of the Alphafold training model, we next utilized the Alphafold 3 server to generate models of KatA_SS1_ in its native tetrameric form, supplying no other information or adjustments to the modeling protocol besides the amino acid sequence. Because we had only four variant sites to use for predictions, and basic chemical knowledge heavily restricts the amount of variability in structural position that would be reasonable at these sites, we limited the number of models we generated to three, which would at least allow us to sample a few different starting seeds and understand the degree to which models might vary. Given the high-quality structural information provided by the four earlier crystal structures, we were unsurprised to find that the global architecture of these Alphafold 3 models mapped well to our new crystal structure (Fig. 2F). However, overlays of these models with our structure also revealed a high degree of accuracy at all four of the variant positions (Fig. 2G-J). Indeed, the models all report high confidence for all regions except for the termini.

After confirming that the AlphaFold 3 server accurately modeled both the global structure of KatA_SS1_ and the variant positions of interest, we investigated whether small changes in user input could affect the results. We generated a new set of models with the following input modifications: (1) single amino acid substitutions at each variant site (with the remainder of the sequence identical to KatA_26695_), (2) insertion of a Trp at either the N-terminus or C-terminus, expected to have minimal impact on global structure or variant positions, and (3) intentional misassignment of the oligomerization state as monomer, dimer, or trimer. To assess similarity in a quantitative way we employed root-mean-square deviation (RMSD) calculations between structural overlays performed across all Cα in a single chain. The two chains of the KatA_SS1_ crystal structure are nearly identical, with an RMSD of 0.14 Å across 488 Cα atoms (Fig. 2K). Comparisons between our KatA_SS1_ crystal structure and AlphaFold 3 models showed RMSD similar to the four KatA_26695_ crystal structures, ranging from 0.27-0.3 Å across 487–488 Cα atoms (Fig. 2K). All models were highly accurate in terms of global RMSD, with the worst, though only by a small margin, being one of the ‘correct’ input models with the true KatA_SS1_ sequence and tetrameric state (Fig. 2K). Hence, small changes to user input, in this case, did not manifest into any substantial changes to global structure.

### Dubious user-input may lower prediction quality for variant sites

Next, we evaluated these models for their ability to accurately predict poses for Val234, Tyr237, Phe255, and Glu421. Overall, the input changes still resulted in poses similar to the KatA_SS1_ crystal structure, but for a couple models the Tyr237 and Glu421 positions had deviations (Fig. 3A-L). For example, of the models with the single position changes, and modeled as the biologically-relevant tetramer, only the Glu421 side chain carboxyl was misplaced (Fig. 3A-D). In the crystal structure the Glu421 O_ε2_ atom forms a hydrogen bond with the guanidinium group of Arg28 from another chain of the tetramer, at a distance of 2.73 Å between the heavy atoms; the shift seen in the AlphaFold3 model is unjustified, moving it to a less favorable 3.14 Å (Fig. 3D). Neither the N- nor C-term Trp additions resulted in any notable decrease in predictive accuracy (Fig. 3E-H).

**Fig. 3.**
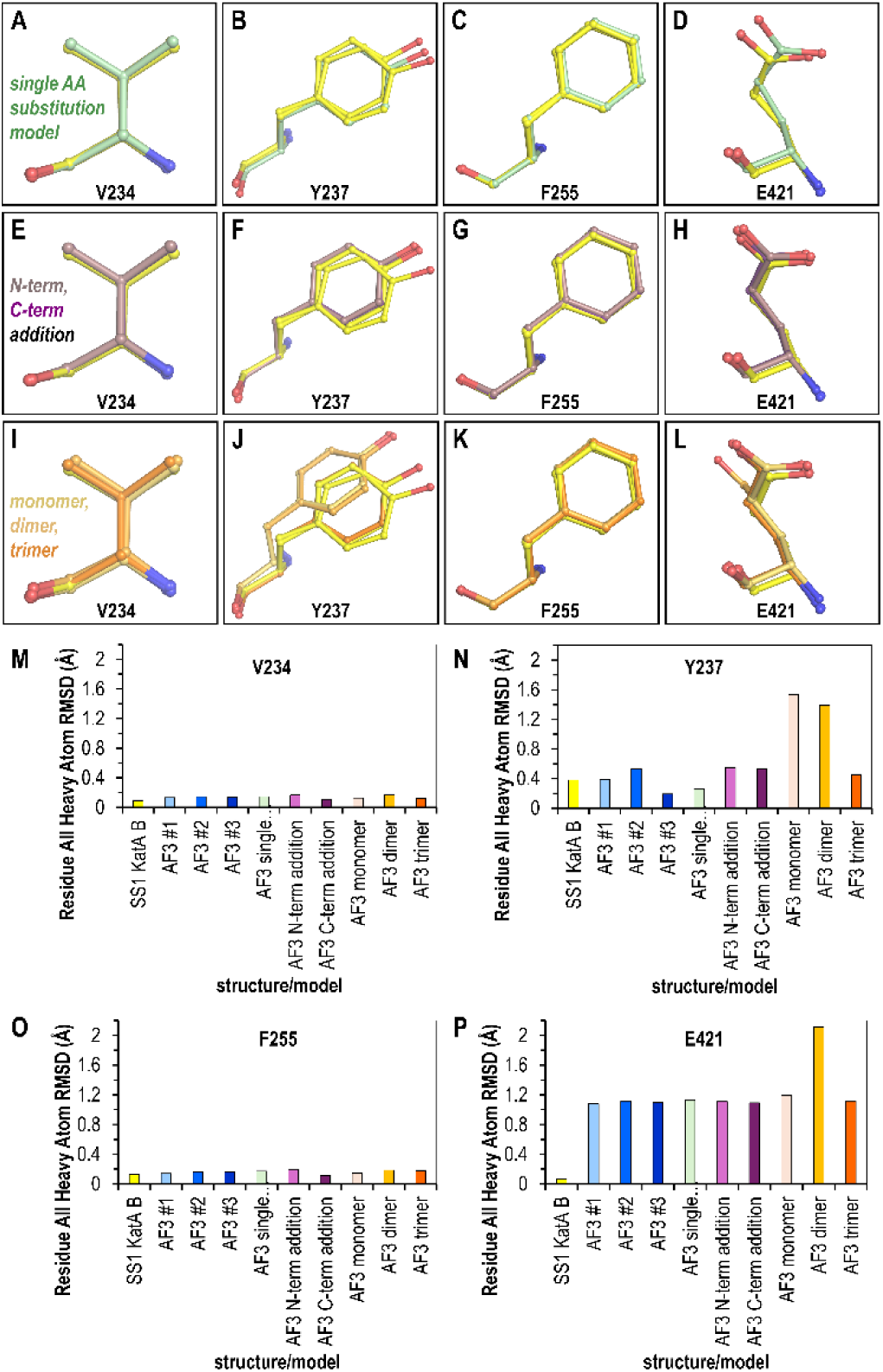
Impact of erroneous user input on Alphafold 3 modeling of variant sites. Overlays show variant sites between the KatA_SS1_ crystal structure (yellow) and Alphafold 3 models generated with: (A–D) single-site mutations (green); (E–H) a single distant Trp insertion at the N-terminus (light purple) or C-terminus (dark purple); and (I–L) the correct sequence modeled with incorrect oligomeric states: monomer (light orange), dimer (orange), and trimer (dark orange). (M–P) Root-mean-square deviation (RMSD) calculations for all heavy atoms compare the Alphafold 3 models to the crystal structure.

However, the set of models most prone to deviation were those with non-native oligomeric states. The Tyr237 position, likely the most challenging site to model, was shifted in the placement of the phenol ring for both the monomer and dimer (Fig. 3J). Since even the experimental structure is poorly resolved at this site, and the residue is solvent-exposed and likely adopts multiple conformations, these deviations may simply reflect that there are multiple energetically-feasible poses. Interestingly, the dimer model contains an entirely different rotamer for the Glu421 position (Fig. 3L). Upon further examination, this could be due to the lack of the Arg28 from the partner chain mentioned above, which would otherwise provide a direct steric clash inhibiting this pose, and the dimer model has made the reasonable guess to shift the Glu421 side chain to be within hydrogen-bonding distance of the nearby His276. Yet, it is unclear why the monomer model, which also is blind to the residues of the tetrameric interface, would not be similarly adjusted, and we presume that this arises from variability in Alphafold 3 outputs.

Examining the RMSD for all heavy atoms between the KatA_SS1_ crystal structure and the Alphafold3 models provides a quantitative comparison for the different strategies (Fig. 3M-P). Across all models, low deviation was seen for the Val234 and Phe255 sites (Fig. 3M, Fig. 3O). Although these sites represent conservative substitutions of similar amino acids, and their structures are arguably the easiest to predict, it is still notable that the deviation from the crystal structure by the Alphafold 3 models is on the same order as the deviation between the two chains within the crystal structure, i.e. these models can be thought of as being at experimental quality (Fig. 3M, Fig. 3O). The Tyr237 site showed the greatest variability in RMSD variation across different models. Striking here is that for some reason the monomer and dimer models, but not the trimer, were the highest deviators, about 2.9-fold greater than other models (Fig. 3N), though there is no clear structural rationale. Lastly, while both chains of the crystal structure have highly similar poses (RMSD of 0.07 Å), the Glu421 site was overall the most challenging for Alphafold 3 to generate highly accurate models, with most near 1.13 Å RMSD, with the dimer, and its different rotamer, the highest at 2.11 Å RMSD (Fig. 3P). Importantly, we note that the AlphaFold model of *H. pylori* KatA_SS1_ currently reported on the UniProt website as part of the ModBase database is a monomer, and shares the incorrect positioning of Glu421 as seen in our AlphaFold 3 monomer model, despite that the per-residue confidence score (predicted local distance difference test, pLDDT) is >90% (AF-F4ZZ52-F1) ^22,34^.

Taken together, the Alphafold 3 models were generally of high quality both globally and at the level of single amino acid prediction for these KatA_SS1_ variants. Variability in model quality does occur even with identical, and correct, user input (Fig. 3M-P). It also seems there may be some risk of lowering model quality if the incorrect oligomeric state is supplied.

### Interpretations of this modeling case study and caveats

In this study, I evaluated how AlphaFold 3 performs in modeling naturally occurring variants of *H. pylori* catalase using only basic input parameters. This system benefits from existing high-quality structural data, likely included in AlphaFold 3’s training set, and the analyzed variants primarily involved conservative amino acid substitutions, presenting relatively straightforward modeling challenges. My key finding is that AlphaFold 3 can produce highly accurate structural predictions of variant residues when provided with the correct sequence and native oligomeric state of the target protein. This is encouraging for researchers without deep expertise in structural biology who rely on AlphaFold 3 for structural insights. However, users should be mindful of specifying the correct oligomeric state. Monomeric models should not be used as proxies for oligomeric structures, as missing tertiary interactions can lead to inaccuracies at the single-residue level.

It is important to note that I used a protein crystal structure as the benchmark for accuracy. Crystallography remains one of the most precise methods for protein structure determination and is well-supported by complementary structural and biophysical approaches ^35–38^. Crystal structures are generally considered reliable representations of proteins in solution and are useful for applications like structure-based drug design ^39–41^. In support of this idea, the various KatA crystal structures, including the novel KatA_SS1_ structure reported here, were solved under different crystallization conditions and crystal forms, yet they maintain a remarkably consistent architecture (Fig. 2A). Crystal structures represent a time- and space-averaged conformation under specific crystallization and data collection conditions ^37,38^. Our structure, like most, was solved under cryocooled conditions, meaning that side chain dynamics at physiological temperatures are likely underrepresented ^42^. Nevertheless, most structures in the Protein Data Bank, used in AlphaFold 3’s training, share these biases ^22,37,38,42^. Without additional experimental evidence, we cannot definitively state whether the deviations observed in our AlphaFold 3 models are biologically relevant, though this possibility remains open (Fig. 3).

Overall, our findings further support the utility of AlphaFold 3 in modeling of protein variants. As the developers and the broader structural biology community continue to focus on improving variant modeling, as reflected in recent studies, this area is poised for further advancement ^4,9,20^. As others have suggested, the most rigorous application of AlphaFold modeling involves using it to test specific hypotheses and interpreting results within the constraints of experimental validation ^18,43^.

## Materials & Methods

### Bioinformatics

The sequence for *H. pylori* KatA SS1 (WP_077231901.1) was retrieved from Uniprot. Geneious was used to perform BLAST searches using standard parameters and retrieve *H. pylori* sequences from the non-redundant amino acid database. Partial sequences were eliminated from further analyses, resulting in 1,931 total sequences, with 1,922 unique (Data S1). MUSCLE ^44^ was used to perform multi-sequence alignment. The resulting aligned sequences were used for analyses of conservation at variant sites of interest and tree-building using Geneious.

### Cloning and protein purification

*H. pylori* KatA from strain SS1 was recombinantly expressed and purified following established protocols with minor modifications ^45^. The *katA* gene was cloned into a pBH vector containing an N-terminal 6×His affinity tag and a TEV protease cleavage site. The construct was transformed into Arctic cells (Agilent) for protein expression. Cells were grown to mid-log phase, induced with 0.4 mM IPTG, and incubated overnight at 15 °C. The protein was purified using nickel affinity chromatography, followed by TEV protease digestion (0.1 mg) to remove the His tag. A reverse His affinity step was performed to isolate the cleaved, untagged KatA. The protein was further purified by size-exclusion chromatography using an S200 column. Enzymatic activity was verified by mixing 10 µl of eluted protein fractions with 100 µl of 30% H_2_O_2_ and 100 µl of Triton X-100, confirming catalase function. The final protein was concentrated in a buffer of 20 mM Tris pH 8, 25 mM NaCl to 10.3 mg/ml and flash-frozen for storage.

### Protein crystallography

Protein crystals were grown using the vapor diffusion hanging drop method. Optimal crystallization conditions for KatA were found to be a 1:1 ratio of protein solution and 0.1 M sodium acetate, pH 4.5, 25% PEG3350. After two weeks of growth at room temperature, large angular crystals appeared suitable for x-ray diffraction analysis. Crystals were incubated with 20% glycerol as a cryoprotectant and then flash frozen in liquid N_2_ for data collection. Diffraction data were collected at the Berkeley Advanced Light Source (ALS) Beamline 5.0.2 with the source at 12398.4 eV. The best-diffracting crystal was chosen for structure determination. Data were processed using DIALS ^46^, and the resolution cutoff set to 1.87 Å. Using a single chain from the crystal structure of PDB: 2a9e the structure was solved through molecular replacement using Phaser ^47^ and found to be in the space group P 2_1_ 2 2_1_ with a dimer in the asymmetric unit, which forms the biologically-relevant tetramer with symmetry mates. To alleviate potential bias in the starting model, the coordinates of the placed solution were randomized by 0.05 Å and the B-factors set to be isotropic and 30 Å^2^. So that the two chains would not be biased to be similar, we did not utilize non-crystallographic symmetry weights. Subsequent model building in Coot ^48^ and refinement with Phenix ^49^, with use of individual B-factors, TLS groups, and riding hydrogens, yielded a final model R/R_free_ of 14.2/19.0%. The final structure has a Molprobity clash score of 2.24, putting it in the 99^th^ percentile for structures of comparable resolution, and uploaded to the PDB as entry 9nh3. Crystallographic statistics are reported in Table S1.

### Alphafold 3 modeling

The Alphafold 3 server at url: https://alphafoldserver.com was used for modeling, using only basic parameters and supplying the sequence of interest and desired oligomer as inputs.

### Structural overlays and RMSD calculations

Overlays and RMSD calculations were performed using the MatchMaker function in ChimeraX. Comparisons of global structure and variant positions were made using overlays in which the entire length of Chain A from crystal structure KatA_SS1_ was used as the reference (across 488 C_α_ atoms).

## Acknowledgments

Funding for this research was provided by NIAID through awards 1K99AI148587 and 4R00AI148587-03, and funding from the College of Veterinary Medicine at Washington State University to AB.

## Author Contributions

A.B. performed the experiments and analyses and wrote the manuscript.

## Supplementary material

**Table S1.** Summary of crystallographic statistics.

| Protein, PDB code | *H. pylori* KatA<sub>SS1</sub>, PDB: 9NH3 |
| --- | --- |
| Space Group | P 2<sub>1</sub> 2 2<sub>1</sub> |
| Cell dimensions and angle<br>(a, b, c, $\alpha$, $\beta$, $\gamma$) (Å, °) | 67.5, 96.2, 154.8, 90, 90, 90 |
| Resolution (Å)<sup>a</sup> | 55.3-1.87 (1.94-1.87) |
| Completeness (%)<sup>a</sup> | 100.0 (100.0) |
| Total reflections | 1087793 (107235) |
| Unique reflections | 84023 (8292) |
| Average $I/\sigma$<sup>a</sup> | 8.45 (1.50) |
| R<sub>merge</sub><sup>a</sup> | 0.28 (1.42) |
| CC<sub>1/2</sub><sup>a</sup> | 0.99 (0.57) |
| R<sub>work</sub> (%) | 14.3 |
| R<sub>free</sub> (%) | 19.0 |
| Ramachandran favored, allowed, outliers (%) | 96.0, 4.0, 0.0 |
| Non-hydrogen atoms | 9435 |
| Solvent atoms | 1205 |
| Protein chains, residues | 2, 978 |
| Average B-factor of protein atoms (Å<sup>2</sup>) | 17 |
| Average B-factor of solvent atoms (Å<sup>2</sup>) | 26 |
| rms bond lengths (Å) | 0.010 |
| rms bond angles (°) | 1.00 |
| TLS groups | 17 |
| Molprobity clash score, percentile | 2.24, 99<sup>th</sup> |

<sup>a</sup> Values in parentheses indicate statistics for the highest resolution shell.

**Data S1.** Sequences of *H. pylori* KatA variants.

